# BAdabouM: a genomic structural variations discovery tool for polymorphism analyses

**DOI:** 10.1101/2020.04.01.018127

**Authors:** Tristan Cumer, François Pompanon, Frédéric Boyer

## Abstract

Genomic Structural Variations (SVs) are known to impact the evolution of genomes and to have consequences on individual’s fitness. Nevertheless, they remain challenging to detect in whole genome re-sequencing (WGS) data. Lots of methods detecting SVs are described in the literature but they might be hard to install, have non-trivial settings, do not detect all SVs categories and have generally high levels of false positive.

Here we introduce BAdabouM, a fast (C written) and easy to install SVs discovery tool. BAdabouM auto evaluates read length, library size and mean coverage to set thresholds specific to each experiment. BAdabouM interprets multiple SVs signatures (reads aligned with a split, non-concordant mapped pairs or uneven coverage) to detect insertions, deletions, copy number variations, inversions, and translocations at single-nucleotide resolution.

When compared with two widely used methods on simulated and real datasets, BAdabouM was faster, exhibited a similar accuracy with a good concordance on SVs detected, and detected significantly more insertions. BAdabouM was more reproducible to detect independently SVs across individuals, which is a clear advantage when characterizing population polymorphism. Furthermore, BAdabouM demonstrated a superior ability to detect breakpoints with a base pair resolution.

BAdabouM proved to be efficient, fast and accurate to detect SVs, and handle. BAdabouM is a complementary method to be used for a more comprehensive detection of SVs, and is especially suited for studying polymorphism for all types of SVs with a high accuracy.

## Introduction

Since decades, population genetics allowed a better understanding of the genetic basis of population differentiation, adaptation, or speciation. Recent rise of Whole Genome Sequencing using massive short paired-end reads sequencing (WGS) gives access to huge amount of Single Nucleotide Polymorphisms (SNPs) among populations increasing phenomenally our knowledge of such evolutionary processes. Those data also contain information about Structural Variations (SVs), which is still underused. SVs are genomic rearrangements generally defined as variations spanning more than 50 bp, including deletions, insertions, inversions, mobile-element transpositions, translocations, tandem repeats, and copy number variations (CNVs) (Tattini, D’Aurizio, & Magi, 2015). SVs possibly have a huge impact on individual’s fitness, inducing phenotypical modifications, diseases susceptibility or local adaptation (Chain & Feulner, 2014). Despite these major roles on individuals and populations, structural variations survey remains uncommon, mainly due to the lack of standardized protocols to reliability detect them in whole genome re-sequencing datasets.

The detection of SVs in WGS data is based on multiple signals. Reads aligned with a split (split reads) allow to detect the breakpoints (i.e. SVs’ start point and end point), while abnormally mapped pairs (e.g., read pairs with both reads aligned with the same orientation or an unlikely long insert size), and uneven depth of coverage indicate the presence of SVs and provide informations to infer their type (Alkan, Coe, & Eichler, 2011).

Several tools are already available to detect SVs (Lin, Smit, Bonnema, Sanchez-Perez, & Ridder, 2014), but they might be hard to install, have non-trivial settings, implies to use arbitrary thresholds and should be run in multiple successive phases lengthening calculation time. Moreover, their joint use on same datasets shows very high false positive rate (Sedlazeck et al., 2018) and low overlap, which justifies a multi tool approach (Pabinger et al., 2014).

## BAdabouM

Here we introduce BAdabouM, a fast (C written), and multi signal integrating tool for discovering structural variations in diploid genomes. BAdabouM self evaluate multiple alignment parameters and use all signals to detect deletion (DEL), insertions (INS), inversions (INV), copy number variation (CNV) and inter and intra-chromosomal translocation (CTX - ITX). BAdabouM detect SVs at a single base pair resolution.

### Input

BAdabouM input file is an indexed bam file, with reads sorted by position.

### Pre-processing and threshold settings

BAdabouM browses part of the file (100k first reads, modifiable option) to auto evaluates experimental characteristics, i.e. read length, library length and mean coverage. Read length and library length are then used to create a sliding window divided in three parts (the first and third ones of the mean size of the library fragments, and the middle one of the mean size of the reads) in order to browse the whole file to detect abnormally mapped reads (respectively pairs of reads) as indicators of SVs. Indeed, BadabouM counts the number of these abnormally mapped reads (respectively pairs of reads) to detect a SV. A key parameter to optimize SVs discovery and avoid false positives is the minimum number of abnormally mapped reads (respectively pairs of reads) that is required to report a SV. The default threshold value, easily modifiable through options, was set to 1/8th of the number of reads in a window for the following reason. In a library-sized sliding window, we expect half of the reads (respectively pairs of reads) to be impacted by a SV in homozygotes and a quarter in heterozygotes. Indeed, half of the reads are aligned forward and half aligned reversed, so at most half of them (or their mate) will be abnormally mapped in case of homozygosity. In the central part of the window, mainly used to the detect breakpoints, we expect all the reads to be splited at the breakpoint in homozygotes and half in heterozygotes. Among the splited reads, half would be on the 5’ side of the breakpoint, and half on the 3’ side (except for deletion or CNV, were the number of reads on each side is not equal). Thus, in heterozygote individuals, a quarter of the reads would be soft-clipped in their 5’ side and a quarter in their 3’ side. This expected number of reads was divided by two to increase the tolerance to coverage variations and take into account a possible uneven representativity of two alleles. Thus, this threshold default value corresponded to half of the expected number of abnormally mapped reads (respectively pairs of reads) in case of heterozygosity.

### Discovery phase

To discover SVs, BAdabouM detects specific signatures of SVs based on split reads, read-pair and depth-of coverage (See supplementary Fig. S1).

An *insertion (INS)* is a region where reads mapped on the forward strand and aligned on the 5’ side of the breakpoint have dangling mate as well as reads mapped on the reverse strand and aligned on the 3’ side of the breakpoint. Reads overlapping the breakpoint must be soft-clipped (i.e., not aligned after the breakpoint or aligned with mismatches after the breakpoint).

A *deletion (DEL)* is a region where pairs of read overlapping the breakpoint have a longer insert size than expected (mean insert size plus two standard deviation, corresponding to the top 5% assuming normal distribution (Pukelsheim, 1994)) and where both breakpoints of the deletion are marked by soft-clipped reads.

A *copy number variation (CNV)* is a region with a higher coverage than expected (2 times the mean coverage) and delimited by two breakpoints highlighted by soft-clipped reads. An *inversion (INV)* is a region with two breakpoints branded by soft -clipped reads, where reads aligned in the forward (respectively reverse) direction 5’ (respectively 3’) of the first (respectively second) breakpoint have their mate orientated the same direction when aligned within the inversion (i.e. between the two breakpoints).

An *intra Chromosomal Translocation (ITX)* is a region where forward reads, aligned before the first breakpoint, have reversed mates mapping on the same chromosome after the second breakpoint of the ITX. Reverse reads, aligned after the first breakpoint, have mates forward mapping before the second breakpoint. All breakpoints are highlighted by soft-clipped reads.

An *inter Chromosomal Translocation (CTX)* is a region where forward reads, aligned before the first breakpoint, have mates mapping on another chromosome on one side of the second breakpoint of the CTX, and where reverse reads after the first breakpoint have mates mapping on the other side of the second breakpoint. All breakpoints are highlighted by soft-clipped reads.

For all types of SVs, the exact location of the breakpoints can be uncertain due to the imprecision inherent to soft-clipped mapping. Thus we report the limits of the range of the breakpoints positions.

### Output

The SVs detected are reported in a table. The first three columns report the chromosome number and the limits of the interval containing the breakpoint position for the beginning of the SV. The three following columns report the same information for the end of the SV. The seventh and eighth columns report the SV type and the length the SV calculated from breakpoint limits. We also provide a script to convert this output to VCF format.

### Expected Results

Considering it’s implementation, BAdabouM’s performances are predictable, with two main limits. The first one is that BAdabouM integrates multiple signals simultaneously to report only high confidence SVs. Thus, BAdabouM may not report true SVs due to the absence of one or more type of signal, such as split read which might be induced by uneven sequence coverage resulting in the absence of reads mapping at a breakpoint. The second limit is due to the use of signals existing only for SVs spanning an area higher than the library size. As a consequence, BAdabouM may not detect deletion, inversions and inter or intra chromosomal translocations smaller than the library size. However, BAdabouM is expected to report high confidence SVs of all types with a single base pair resolution of the breakpoint.

## Application

To test BAdabouM’s ability to detect SVs in both simulated and real datasets, we compared it with two commonly used methods, Delly (Rausch et al., 2012) and Breakdancer (K. Chen et al., 2009). Theses two softwares were selected for their ability to detect a wide range of SVs. Delly detects deletions, inversions, duplication and inter- and intra chromosomal translocations based on abnormally mapped pairs, while Breakdancer detects deletions, inversions, duplication and insertions based on both abnormally mapped pairs and split read). They were also chosen because they are widely used by the scientific community to analyse datasets alone (Zhao et al., 2016) or combined in pipelines (Mimori et al., 2013; Mohiyuddin et al., 2015), and also to benchmark new softwares (Layer, Chiang, Quinlan, & Hall, 2014; R. Chen, Lau, Zhang, & Yang, 2016; Chong et al., 2017)). We may note that Delly does not detect Insertion and BreakDancer does not detect CNVs.

### Simulated dataset

Softwares were first benchmarked using a set of simulated genomes harbouring each types of structural variant. This panel was composed to test the detection of Insertions, Deletions, Inversions, Copy Number Variations, and Inter and Intra-chromosomal translocation. For each type of SVs, 25 events distributed over five different sizes were simulated (five event of each size: 100, 250, 500, 1000 and 5000 bp). These SVs were simulated in the first 1M bp of the first two chromosomes of the sheep reference genome OAR_v4, after removal of undefined nucleotides, i.e. N).

We then simulated resequencing data for one hundred homozygous and one hundred heterozygous individuals, with a sequencing coverage of 20X and a library size of 300bp (Huang, Li, Myers, & Marth, 2012). More details about simulated data are available in supplementary informations. Software were then evaluated based on (i) their ability to detect the event and (ii) the precision of the detection (distance between predicted and simulated breakpoints).

The results of the simulations (Table 1) show that softwares performances vary across SVs types and size. Considering SVs discovery rates, the performances of BAdabouM are similar to that of the two other softwares, except a lower discovery rate for inversions. Alike, BadabouM had a lower ability to detect SVs at the heterozygous state than the two other methods, inducing higher false negatives rate (Table 1). It’s interesting to note that BAdabouM was able to detect insertions much better than Breakdancer (Delly does not detect insertions). The softwares were clearly different in their precision in predicting breakpoints. BAdabouM located SVs with very high accuracy (median distance between predicted and real breakpoint ranging from 1 to 1.5bp, except for 250bp inversion where breakpoint precision was 28bp), while Delly was less accurate (median distance ranging from 14bp to 60bp) and breakdancer was even less precise (median distance ranging from 16bp to 286bp).

**Table 1:** Benchmarking of three SVs discovery tools on simulated datasets. For each type of SVs, the first value corresponds to the median detection rate over five replicates; the value between brackets corresponds to the median distance between estimated and simulated breakpoints.

| Type | Size | BAdabouM |  |  |  | Delly |  |  |  | BreakDancer |  |  |  |
| --- | --- | --- | --- | --- | --- | --- | --- | --- | --- | --- | --- | --- | --- |
|  |  | Homozygote |  | Hetrozygote |  | Homozygote |  | Hetrozygote |  | Homozygote |  | Hetrozygote |  |
|  |  | detec. rate | precision | detec. rate | precision | detec. rate | precision | detec. rate | precision | detec. rate | precision | detec. rate | precision |
| INS | 100 | **49** | **(1)** | **6** | **(1)** |  |  |  |  | 1 | (28) | 0 |  |
|  | 250 | **96** | **(1)** | **85** | **(1)** |  |  |  |  | 18 | (22.75) | 4 | (22.25) |
|  | 500 | **97** | **(1)** | **85** | **(1)** |  |  |  |  | 0 |  | 0 |  |
|  | 1000 | **88** | **(1)** | **48** | **(1)** |  |  |  |  | 0 |  | 0 |  |
|  | 5000 | **98** | **(1)** | **82** | **(1)** |  |  |  |  | 0 |  | 0 |  |
| DEL | 100 | 0 |  | 0 |  | 0 |  | 0 |  | **5** | **(98)** | **2** | **(87.25)** |
|  | 250 | 0 |  | 0 |  | 40 | (60) | 28 | (60.75) | **100** | **(19.5)** | **99** | **(16)** |
|  | 500 | **100** | **(1.5)** | 98 | **(1.5)** | **100** | (17) | **100** | (15) | **100** | (20.5) | **100** | (17.5) |
|  | 1000 | **100** | **(1)** | 99 | **(1)** | **100** | (17.5) | **100** | (15.25) | **100** | (20.5) | **99** | (18.5) |
|  | 5000 | **100** | **(1.5)** | 98 | **(1.5)** | **100** | (16.5) | **100** | (15.25) | **100** | (21) | **100** | (18.25) |
| CNV | 100 | 0 |  | 0 |  | 0 |  | 0 |  |  |  |  |  |
|  | 250 | 95 | (27.375) | **95** | (28) | **96** | **(16.25)** | 80 | **(15.25)** |  |  |  |  |
|  | 500 | 99 | **(1)** | 98 | **(1)** | **100** | (19.5) | **100** | (18.5) |  |  |  |  |
|  | 1000 | 98 | **(1.5)** | 97 | **(1.5)** | **100** | (20.5) | **100** | (19) |  |  |  |  |
|  | 5000 | 99 | **(1)** | 99 | **(1)** | **100** | (19.5) | **100** | (18.5) |  |  |  |  |
| INV | 100 | 0 |  | 0 |  | **100** | **(24)** | **99** | (32.5) | **100** | (156) | **99** | (119.5) |
|  | 250 | 0 |  | 0 |  | **100** | **(15.75)** | **100** | **(15.125)** | **100** | (185.75) | **100** | (153) |
|  | 500 | 68 | **(1.25)** | 30 | **(1.25)** | 100 | (16) | 100 | (14) | 100 | (189) | 100 | (172.125) |
|  | 1000 | 74 | **(1.25)** | 32 | **(1.25)** | 100 | (15.5) | 100 | (14.25) | 100 | (280.25) | 100 | (236) |
|  | 5000 | 72 | **(1)** | 35 | **(1)** | 100 | (15.5) | 100 | (13.5) | 100 | (277.75) | 100 | (244.5) |
| ITX | 100 | 0 |  | 0 |  | 0 |  | 0 |  | **92** | (27) | **89** | (46) |
|  | 250 | 0 |  | 0 |  | 0 |  | 0 |  | **3** | (183) | **27** | (80) |
|  | 500 | **99** | **(1)** | **76** | **(1)** | 0 |  | 0 |  | 10 | (274) | 53 | (235) |
|  | 1000 | 99 | **(1)** | 76 | **(1)** | 0 |  | 0 |  | **100** | (279.5) | **98** | (244.5) |
|  | 5000 | 99 | **(1)** | 77 | **(1)** | 0 |  | 0 |  | **100** | (280) | **97** | (242.5) |
| CTX | 100 | 0 |  | 0 |  | **100** | (14.75) | 92 | (25.25) | **100** | (129) | **97** | (105) |
|  | 250 | 0 |  | 0 |  | **100** | (16) | **99** | (20.375) | **100** | (172.25) | **99** | (119.125) |
|  | 500 | 96 | **(1)** | 77 | **(1)** | **100** | (16.5) | **100** | (15) | **100** | (278) | **100** | (253.5) |
|  | 1000 | 88 | **(1)** | 71 | **(1)** | **100** | (16.5) | **100** | (14.5) | **100** | (286.5) | **100** | (255.5) |
|  | 5000 | 88 | **(1)** | 71 | **(1)** | **100** | (16.25) | **100** | (15) | **100** | (272.5) | **100** | (239) |

Combining discovery rate and precision, BAdabouM could detect all types of structural variations at a single base pair resolution. The lower detection rate observed (Table 1) is probably due to the fact that BAdabouM integrates multiple signals simultaneously. The cost to pay for this precision is a higher rate of false negatives. Combining BAdabouM with other softwares would maximise accuracy for both SV location and detection success. We can note that none of the softwares detected false positives in simulated data.

### Real dataset

To benchmark the different methods on real data, we took advantage of a recently published dataset of medium coverage whole genome sequences (about 12X) from 53 individuals from wild and domestic sheep (three *Ovis* species)(Alberto et al., 2018, Supplementary table 1), The goals were to (i) test BAdabouM’s ability to discover SVs on real data, (ii) examine the concordances between BadabouM and the two other methods and (iii) compare the computational resources needed by the three softwares.

A correspondence analysis (Fig. 1A) showed that BadabouM was able, as the two other methods, to detect the genetic signals that fully differentiate the *Ovis* species. While 11,5 % of the whole set of variants discovered were detected by all three methods, a greater part (32,5 %) was shared only by two methods, and the majority (56 %) was specific to one software (Fig. 1B). This low concordance between methods highlights the technical difficulty to detect SVs, a general concern that has already been emphasised (Pabinger et al., 2014). However, we must consider that 80% of the events detected by BAdabouM alone were insertions that are not detected by Delly, and for which Breakdancer performs poorly as shown with the simulated dataset. Thus, if we except insertion, a high concordance of 91.1% was observed between BAdabouM and the two other softwares.

**Figure 1:**
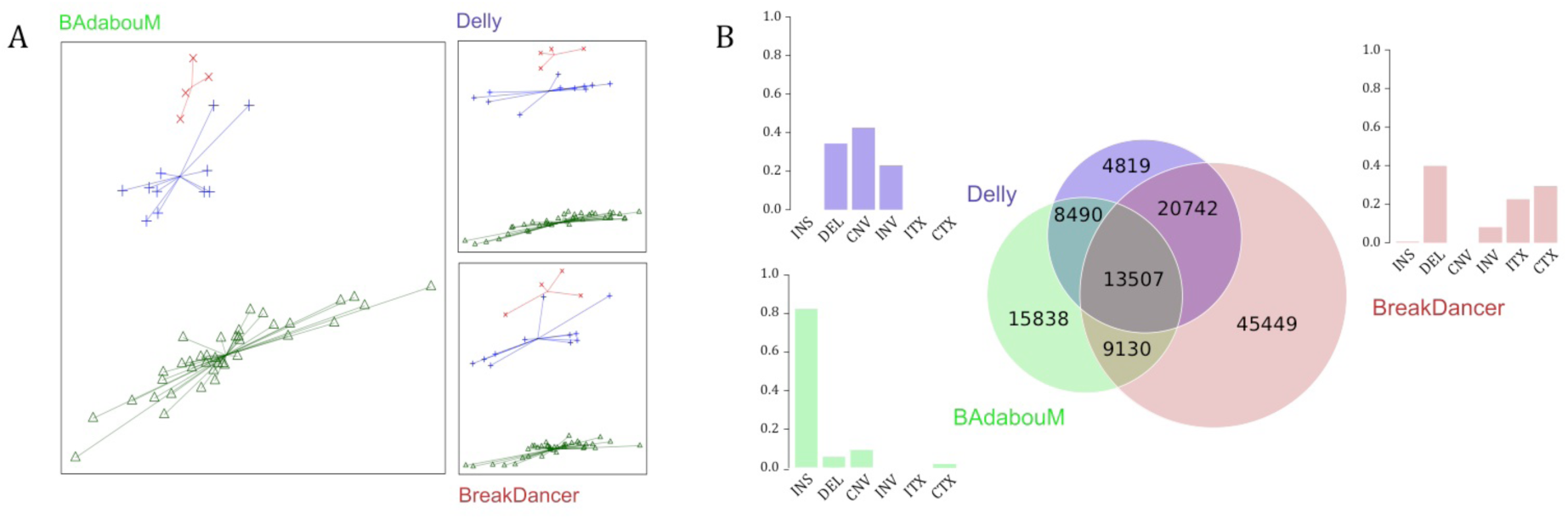
Comparison of SV detection by BAdabouM, Delly and Breakdancer on a real dataset of 53 sheep WGS. (A) First two axes of a correspondence analysis based on the SVs discovered by each software. Each point corresponds to an individual from an *Ovis* species: Urial, (*Ovis vignei, red “x” symbol)*, Asiatic mouflon (*Ovis orientalis, blue plus symbol)* and domestic sheep (*Ovis aries*, green triangles). (B) Venn diagram of predicted SVs by each software. Barplots summarize the proportion of each type of SVs specifically detected by each software.

Interestingly, variants detected by BAdabouM in an individual were more frequently detected independently in another individual (i.e., overlap Table 2) than with the two other methods. The lower number of SVs globally detected and the higher overlap between individuals reflects the BAdabouM’s stringency, which relies on the simultaneous detection of multiple signals for high confidence SVs. Moreover, the measurement of computation time (Table 2) highlighted that BadabouM performed noticeably faster.

**Table 2:** Software performances for running time, mean number of structural variation per individuals and proportion of SVs discovered independently in at least two individuals.

| Software | time (min) | nb of SVs / indiv | Overlap (%) |
| --- | --- | --- | --- |
| BAdabouM | 44 | 5607 | 95.8 |
| Breakdancer | 55 | 16740 | 92.1 |
| Delly | 165 | 7384 | 91.3 |

## Software originality

BAdabouM is easy to install and handle, fast, and can detect all types of structural variations in whole genome re-sequencing data. Moreover, it is more conservative than other softwares by integrating all signals to detect SVs for maximising the avoidance of false positives. It also performs better to detect a given SV across individuals with a very high accuracy at single base pair resolution. All these properties are especially suited for the characterization of SVs polymorphisms in population genomics approaches (Luikart, England, Tallmon, Jordan, & Taberlet, 2003) aiming at characterizing thousands of variants in dozens to hundreds of individuals based on genome scans or re-sequencing data.

## Supporting information

Supplementary_InformationsAndTables

## Acknowledgements

This work was supported by the Labex Osug@2020 (Investissements d’avenir — ANR10LABX56)

All computations presented in this paper were performed using the CIMENT infrastructure (https://ciment.ujf-grenoble.fr), which is supported by the Rhône-Alpes region (GRANT CPER07_13 CIRA: http://www.ci-ra.org) and France-Grille (http://www.france-grilles.fr).

## Authors’ contributions

TC and FB designed and implemented the software. All authors contributed to the design of the analyses conducted by TC. TC wrote the manuscript with the help of FB and FP.

## Data Accessibility

Genome sequences used in this work are available at http://projects.ensembl.org/nextgen.

BAdabouM is distributed under the CeCILL license and is freely available at http://github.com/cumtr/badaboum.

