## Supplementary_InformationsAndTables for "BAdabouM: a genomic structural variations discovery tool for polymorphism analyses": BAdabouM_Supplementary_file_JoHe.pdf

<u>Supplementary informations:</u>

### 1. BAdabouM

#### *1.1 Detection of Structural Variations*

BAdabouM implements a window sliding along the genome for detecting specific signatures in paired-reads mapping. This sliding window is divided in three parts, allowing to detect, on its both sides, abnormally mapped reads and uneven coverage, and split reads in the middle one (Figure S1).

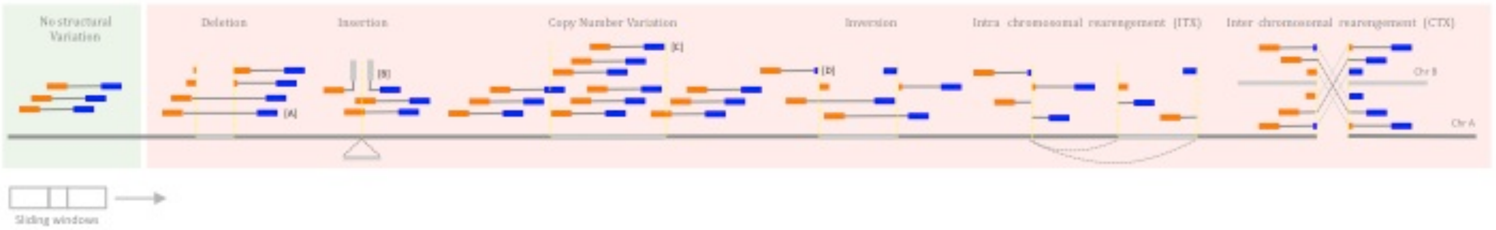


Figure S1 : schematic representation of the sliding windows and specific signatures of different types of SVs. To discover SVs, BAdabouM detects inconsistent alignments: [A] A space between pairs of reads.[B] Pairs of reads with one read not mapped close to its pair (i.e., dangling).[C] Too many or too few reads are aligned onto the sequence.[D] Split-reads confirm the occurrence of a SV and allow precise location of the breakpoints.

### 2. Application – Simulated dataset

#### *2.1 Simulations*

The simulated panel of SVs was composed of Insertions, Deletions, Inversions, Copy Number Variations, and Inter and Intra-chromosomal translocations. An insertion is a sequence that is not present in the reference but present in the individual genome. A deletion is a sequence that is not in the individual genome but in the reference sequence. A CNV is a sequence present once in the reference, and five times (consecutives) in the simulations. Inter and intra chromosomal translocations are sequences that are in both the individual genome data and the reference sequence, but at different locations. For each type of SVs, five sizes were simulated (100, 250, 500, 1000 and 5000 bp), with five replicates for each size of SVs. A part of the sequence of chromosome 1 and 2 of the sheep genome (Oar_v4.0, https://www.ncbi.nlm.nih.gov/assembly/GCF_000298735.2), with Ns removed, was modified according to the tested SV (see Supplementary tables 2 and 3 for complete description of the sequences used to simulate SVs).
Simulated sequences are available under request. Sequencing data were then simulated for one hundred homozygous and one hundred heterozygous individuals using the ART software (defaults parameters, except coverage: 20X and library size 300bp) (Huang, Li, Myers, & Marth, 2012).

#### *2.2 SVs discovery*

SVs where detected independently for each simulated individual using three different methods; BAdabouM, Delly (Rausch et al., 2012) and Breakdancer (Chen et al., 2009). All three methods were run using default parameters, except for the mapping quality, where a quality of 60 was required.

*2.3 Simulation summary*

Estimation of SV calling performances: for each software we estimated the detection rate of each variant and the distance between the estimated and simulated breakpoints.

### 3. Application – real dataset

*3.1 Sequences origin*

The paired-end raw resequencing data of the genomes of the 53 individuals from three *Ovis* species (*O. vignei*, *O. orientalis* and *O. aries*) used in (Alberto et al., 2018, also described in Supplementary table 1) were downloaded from http://projects.ensembl.org/nextgen.

*3.2 Alignment*

100bp paired-end reads for *Ovis* were mapped to the sheep reference genome (build Oar_v4.0 - GenBank assembly accession: GCA_000298735.2) using BWA-MEM (Li & Durbin, 2009). The BAM file produced for each individual was sorted using Picard SortSam and improved using Picard MarkDuplicates (http://picard.sourceforge.net).

*3.3 SVs calling*

SVs were called independently for each individual using three different methods; BAdabouM, Delly (Rausch et al., 2012) and Breakdancer (Chen *et al.*, 2009). All three methods were run using default parameters, except for the mapping quality, where a quality of 60 was required.

*3.4 SVs merging and filtering*

For each individual, SVs detected by two methods were considered as identical based on the reciprocal overlap of their position. As methods do not detect breakpoints with the same accuracy, a threshold was set for congruent overlap. For inversions, deletions, duplications and intra-chromosomal translocations, a criterion of SV overlap greater than 50% was used. For insertion, as breakpoints may be close, SVs were considered as identical once an overlap existed whatever its size. For intra and inter-chromosomal translocations, breakpoints within a 1kb window were considered as corresponding to the same event.

3.5 Analysis

All statistics were run using the R language (Ihaka & Gentleman, 1996). AFC were run using the ade4 package (Dray et al., 2017, p. 4).
